# The role of Aβ circRNA in Alzheimer’s disease: alternative mechanism of Aβ biogenesis from Aβ circRNA translation

**DOI:** 10.1101/260968

**Authors:** Dingding Mo, Xinping Li, Carsten A. Raabe, Timofey S. Rozhdestvensky, Boris V. Skryabin, Juergen Brosius

## Abstract

Alzheimer’s disease (AD) is an age-related detrimental dementia. Amyloid beta peptides (Aβ) play a crucial role in the pathology of AD. In familial AD, Aβ is generated from the full-length amyloid precursor protein (APP) *via* dysregulated proteolytic processing. However, Aβ biogenesis in case of sporadic AD remains so far elusive. circRNAs are a class of transcripts, which are preferentially expressed in brain. Here, we identified a circRNA (circAβ-a) derived from the Aβ-coding region of the *APP* gene. circAβ-a is expressed in brains of AD and nondementia controls. With the aid of our recently established approach for analysis of circRNA functions, we demonstrated that circAβ-a is efficiently translated into a novel Aβ-related Aβ175 protein in living cells. Importantly, Aβ175 was further processed to Aβ peptides, the hallmark of AD. In summary, our analysis revealed alternative route of Aβ biogenesis. circAβ-a and its corresponding protein might represent novel therapeutic targets for AD treatment.

## INTRODUCTION

Alzheimer’s disease (AD) is one of the most common and devastating forms of dementia. It is associated with the gradual loss of cognitive abilities such as memory and thinking; the progression of AD also is accompanied by various behavioural issues [1–4]. Naturally, as an age-related, neurodegenerative disorder, the greatest known risk factor is increasing age itself [3,5]. The pathogenesis and clinical manifestation of familial AD is accompanied by the formation of insoluble amyloid beta (Aβ) plaques and neurofibrillary tangles (NFTs) [1–4,6–10]. Although there are still debates about the actual role of Aβ in AD pathogenesis[11], extensive human genetics studies confirmed its strong association with AD disease development[1,12,13]. Aβ peptides are generated from the full-length amyloid precursor protein (APP) *via* sequential proteolytic processing by beta (β) and gamma (γ) secretases [13,14]. Initially, the N-terminal domain of APP is cleaved by β-secretase, subsequentially γ-secretase cleaves the remaining C-terminal fragment of APP (CTF99) to generate Aβ [14]. Dysregulation of this event is believed to be responsible for Aβ accumulation [14]. Several mutations within the CDS (protein coding sequence) of APP were identified and the specific association of these genetic variants with increased accumulation of Aβ were established [12,15–23]. Similarly, some diagnostic mutations within presenilin genes, which encode the catalytic subunits of the gamma secretase complex, are linked to increased Aβ levels [24–32]. Subsequent polymerization of Aβ peptides leads to oligomers, which in turn aggregate to insoluble amyloid plaques. This process is believed to cause tau hyperphosphorylation. The resulting formation of neurofibrillary tangles initiates a complex cascade of cellular reactions that ultimately are leading to neuronal death [1,2,23,26,29,33,34].

However, this pathogenesis is common only to familial AD, which accounts for 1–5 % of all cases worldwide [1,2]. In fact, the most common form of AD is the “sporadic” variant. This specific form of AD prevails in patients aged 65 or older and unlike familial AD, no specific mutations in APP and presenilin genes were identified. However, apart from these striking genetic differences both forms are characterized by the overproduction of Aβ plaques in brain [2,11,12,14,35,36]. Moreover, although beta-secretase (BACE1) expression and its enzymatic activity were reported to increase in most sporadic AD patients, APP full length protein and γ-secretase proteolytic activity remains unchanged (compared to non-demented controls) [37,38]. In addition, extensive analysis of transgenic mouse models for monitoring pathophysiological consequences of human wildtype (i.e., unmutated) APP overexpression did not reveal signs of increased Aβ plaques formation in brain [39–42]. Hence, the underlying mechanism of Aβ production in case of sporadic AD remains widely elusive. The *in vivo* data, however, might suggest that there is potentially an alternative route and/or precursor leading to the generation of Aβ peptides.

Circular RNAs (circRNAs) are abundant products from transcripts[43]; mostly derived from protein coding genes[44,45]. They are the result of hnRNA/pre-mRNA back-splicing and as such represent covalently closed circles, which are devoid of RNA cap structures or terminal poly(A) tails, features, which render them substantially different from their corresponding linear mRNA counterparts [44,46]. Previous analysis revealed that circRNA formation is most prominent in brain [47–50]. Interestingly, in several organisms, circRNAs are being regulated in an age-dependent manner. This might suggest that circRNAs play regulatory roles during ageing and potentially even are responsible for the development of age-associated neurodegenerative diseases [46,49,51–54]. More importantly, recent studies demonstrate that certain circRNAs encode proteins, suggesting that they might play essential biological roles similarly to linear mRNAs [55–58].

We detected a circRNA derived from the *APP* gene in human brain samples of AD patients and non-dementia controls. Since the circRNA contains the corresponding Aβ coding sequence, it is referred to as circAβ-a. Recently, we exploited ‘intron-mediated enhancement’ (IME), and successfully established a method for the investigation of circRNA translation in living cells [55]. We demonstrated that circAβ-a serves as template for synthesis of a novel Aβ-related protein variant. Notably, the resulting product is further processed, leading to Aβ peptides in human HEK293 cells. Our findings, therefore, established an alternative route to Aβ biogenesis in human cells via circRNA translation expressing a C-terminal polypeptide variant, the canonical mRNA cannot generate.

## RESULTS

### Presence of circAβ-a in the brain of AD patients and non-dementia controls

Mechanisms of Aβ production in sporadic AD remain elusive. In search of alternative routes to Aβ biogenesis we analysed circRNAs, which are derived from the Aβ coding region. circRNA hsa_circ_0007556 harbours APP protein coding exons 14 to 17, is 524 nt in length and has been detected in several independent RNA high-throughput sequencing datasets [43,47,59,60]. Here we addressed whether a potential protein product of hsa_circ_0007556 (henceforth referred to as circAβ-a) can constitute an alternative route to Aβ peptide generation. For detection of circAβ-a in various human brain samples, we performed RT-PCR with two pairs of divergent primers. To account for the interindividual variance of circAβ-a levels, we included six independent samples (prefrontal cortex total RNA) representing AD patients (three) and non-dementia controls (three), respectively (Fig. 1A). In our analysis, reverse primer circAβ-a-R1 mapped to the splice junction region of circAβ-a; this design ensured the specificity of our PCR reaction (Fig. 1A). In addition, due to the divergent (relative) orientation of our primer pair, this PCR assay required circular molecules as a template. We only detected a single amplification product of the expected size (499 bp), i.e. for all AD and non-dementia control samples (Fig. 1B). For further increase of specificity, a RNase R pre-treatment of total RNA samples was incorporated, which enables the digestion of linear RNA but leaves circular molecules unaffected. As anticipated, we uncovered the same size (499 bp) amplification product. Sanger sequencing of the resulting PCR amplicon confirmed that the circRNA contained only APP protein coding exons 14-17 and no intronic sequences were included (Supplementary data-1). These results were confirmed with a second pair of oligonucleotide primers (circAβ-a-F2 and circAβ-a-R2, Fig. 1A and B). Migration of the resulted amplicons in agarose gel exhibited PCR products of ~150 bp, in agreement with the expected amplicon length of 148 bp. Again, the same product was obtained by RNase R pre-treatment prior to RT-PCR assays, once more confirming the circular structure of circAβ-a. Sanger sequencing of the corresponding RT-PCR product confirmed the anticipated sequence of circAβ-a (Supplementary data-2). The resulting circAβ-a junction sequence is displayed in Fig. 1C. In summary, we demonstrated the presence of circAβ-a in the human brain, both in AD and non-dementia samples by RT-PCR and sequencing.

**Fig. 1.**
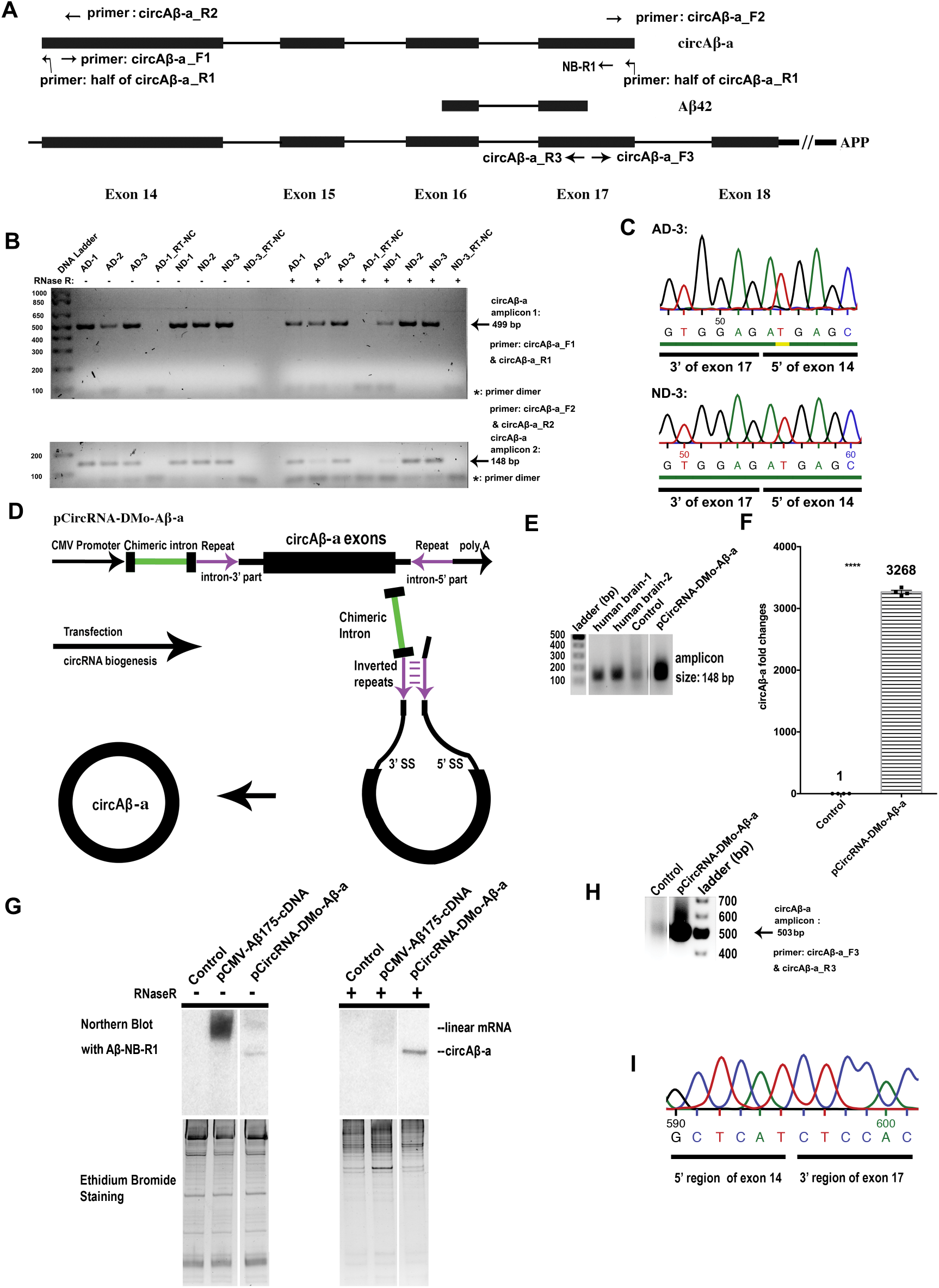
Identification of circAβ-a formation in the human brain and its transient overexpression in HEK293 cells. **A. Position of circAβ-a within the APP coding region.** The Aβ42 corresponding sequence is indicated as reference and maps to protein coding exons 16 and 17. circAβ-a consists of exons 14, 15, 16, 17. Primers for RT-PCR and Northern blot hybridization are indicated. **B. Agarose gel electrophoresis (1%) of circAβ-a RT-PCR products.** Amplifications were performed with a set of divergent oligonucleotide primers (circAβ-a_F1 and circAβ-a_R1, see upper panel) on human prefrontal cortex total RNA from AD and non-dementia controls as template. Primer pair circAβ-a_F2 and circAβ-a_R2 was employed for circAβ-a amplification by qRT-PCR; the resulting PCR products were also analysed by 1% agarose gel electrophoresis (see lower panel). An RNase R digestion step was incorporated for the enrichment of circular RNA. RT-NC: the negative control for reverse transcription, i.e. samples treated devoid of reverse transcriptase. **C. Sequencing of RT-PCR products (see B for details), results demonstrate the fusion of exon 17 (3’ end) to exon 14 (5’ end), representing the actual junction region of circAβ-a.** cDNAs were generated with RNase R treated total RNA of AD-3 and ND-3 samples with divergent primer pair (circAβ-a_F2 and circAβ-a_R2). **D. circAβ-a overexpression constructs.** Figure D is modified from [55]. Our circRNA formation strategy based on intron-mediated enhancement (IME) is schematically depicted **E. RT-PCR amplification confirmed circAβ-a formation in human brain samples and recombinant HEK293 cells overexpressing circAβ-a.** Control represents mock transfection with pCircRNA-DMo empty vector, pCircRNA-DMo-Aβ-a depicts circAβ-a formation after transfection with pCircRNA-DMo-Aβ-a (i.e. the circAβ-a expression construct). **F. Quantification of circAβ-a levels in HEK293 cells.** Control represents mock transfection with empty vector (pCircRNA-DMo), Fold changes (y-axis) and Student’s T-tests were performed in comparison to the control sample, ****, P ≤ 0.0001, n = 4. Mean ± SEM of Control: 1.019 ± 0.1216, n=4; Mean ± SEM of pCircRNA-DMo-Aβ-a: 3268 ± 27.22, n=4. **G. Northern blot analysis of circAβ-a formation in HEK293 cells.** The pCMV-Aβ175-cDNA expression vector harbours the linear ORF corresponding to circAβ-a under the control of the CMV promoter for expression. - depicts untreated samples; + depicts RNase R treated samples, ethidium bromide staining of the 15 μg total RNA in native 5% polyacrylamide gels served as the loading control. **H. Agarose gel electrophoresis (1%) for circAβ-a RT-PCR products (503 bp)**. RT-PCR reactions were conducted with primers circAβ-a_F3 and circAβ-a_R3. **I. Sanger sequencing for the Aβ junction region**. PCR products (see above) were cloned and sequenced showing the circAβ-a junction region representing RT-PCR products from the pCircRNA-DMo-Aβ-a transfection. The results also confirmed that circAβ-a was identical to spliced CDS exons 14, 15, 16 and 17 of the *APP* gene (data not shown).

### Intron-mediated enhancement (IME) promotes circAβ-a overexpression in human cells

We recently established a novel system for the analysis of putative circRNA functions based on intron mediated enhancement [55]. This approach helps to efficiently express circRNAs in transient transfection assays and to investigate mechanisms of circRNA translation. We adapted this universal method to analyze the putative protein coding ability of circAβ-a. For this purpose, a cDNA representing APP protein coding exons 14-17 was inserted into pCircRNA-DMo plasmid, which harbours a chimeric intron to enhance circRNA formation (Fig. 1D) [55]. In transiently transfected HEK293 cells, the resulting construct (pCircRNA-DMo-Aβ-a) demonstrated robust circAβ-a levels (Fig. 1E). Importantly, agarose gel electrophoresis of RT-PCR products confirmed that the product of heterologous circAβ-a formation in HEK293 cells and the endogenous transcript in the human brain are of identical size (Fig. 1A and E). Analysis of the total RNA by qRT-PCR revealed 3268-fold overexpression of circAβ-a, compared to the endogenous background levels in HEK293 cells (Fig. 1F).

For further validation, we also resorted to RNase R treatment of total RNA followed by Northern blot hybridization. This experimental design required controls for RNase R digestion and Northern hybridization. For this purpose, we generated vector pCMV-Aβ175-cDNA, which enables the expression of the linear circAβ-a RNA variant including the C-terminal non-canonical amino acid residues from the circular template in control transfections. Two specific signals with different migration in 5% native polyacrylamide gels were discriminated between circRNA and linear mRNA by Northern blots following transfection. The slower migrating RNA in pCMV-Aβ175-cDNA transfections did not remain detectable after RNase R treatment (Fig. 1G), suggesting that the signal corresponds to the linear variant of circAβ-a. On the other hand, the faster migrating RNA in circAβ-a transfections (pCircRNA-DMo-Aβ-a) - was resistant to RNase R digestion, which is in agreement with its portrayed circular structure (Fig. 1G). Similar to our previous report [55], no linear RNA variants were detectable by Northern blot hybridizations in pCircRNA-DMo-Aβ-a transfections, in turn minimizing chances of linear RNA ‘contamination’ in subsequent assays (see below).

By RT-PCR amplification and subsequent Sanger sequencing with a third pair of oligonucleotide primers (circAβ-a_F3 and circAβ-a_R3, Fig.1A), we were also able to reconfirm the correct splice junction of circAβ-a in HEK 293 cell transfection experiments (Fig. 1H and I). In addition, the qRT-qPCR analysis of APP mRNA revealed that the endogenous APP mRNA levels remained largely unaffected (Supplementary data-3). We concluded that our IME strategy enabled strong circAβ-a formation in HEK293 cells. Notably, HEK293 cells represent a well-established cellular model for Alzheimer’s disease [61,62]; our system is therefore appropriate for the *ex vivo* analysis of potential circAβ-a functions in AD.

### circAβ-a can be translated into an Aβ-related protein in human cells

Recent reports demonstrated that open reading frames (ORFs) of circular RNAs can act as templates for efficient protein biosynthesis [55,56,58]. We identified a hypothetical 528 nt long ORF, presumably beginning at the first potential AUG start codon in exon 14 and ending, after what corresponds to the 3’ end of exon 17, out of frame into what corresponds to the 5’ end of exon 14. This adds 17 codons to the C-terminus of the circAβ-a sequence (see below). This translates into a putative polypeptide of maximally 175 amino acids (aa) of approximately 19.2 kDa (Fig. 2A and B, Supplementary data-4). This circAβ-a-derived protein is henceforth referred to as Aβ175 (Fig. 2B). To investigate whether circAβ-a encodes a *bona fide* protein, we evaluated Aβ175 expression in transient transfection assays of pCircRNA-DMo-Aβ-a in HEK293 cells by Western blot analysis. In order to avoid proteolytic processing of Aβ175, several secretase inhibitors were added to the cell culture medium (Materials and Methods). Plasmid pCMV-Aβ175-cDNA, expressing the linear mRNA form of circAβ-a ORF (see above), was utilized in parallel transfection assays and served as positive control. Western blot analysis with the amyloid beta peptide antibody (WO2) revealed that the circAβ-a ORF is efficiently translated into an Aβ-related protein of approximately 25 kDa (between the 22 kDa and 36 kDa size markers) (Fig. 2C). Notably, signals for the circAβ-a-related protein and the positive control (pCMV-Aβ175-cDNA) were of identical size.

**Fig. 2.**
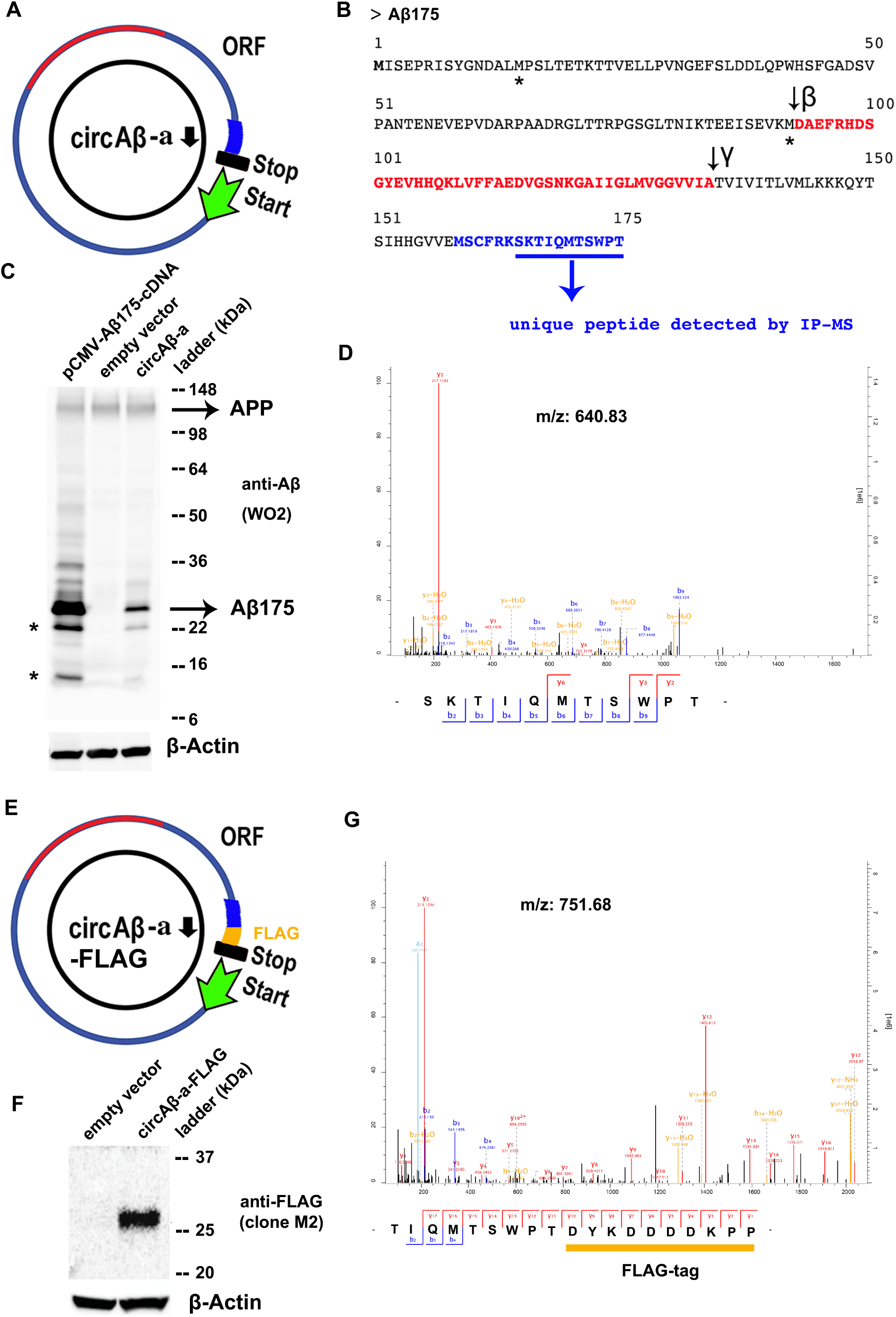
Analysis of circAβ-a translation and Aβ-related protein (Aβ175) in HEK293 cells. **A. The open reading frame (ORF) of circAβ-a is represented by a blue circle.** The bright blue colour depicts the unique peptide sequence of the circAβ-a translated protein, the green arrow indicates the presumed translation start codon, the black bar represents the translational stop codon; the inner black arrow indicates the 5’ nucleotide of exon 14. **B. The polypeptide sequence of Aβ175.** Black and red letters represent the polypeptide sequence of Aβ175, which is identical with the corresponding segment of wild type APP protein. Processing sites for β and γ secretases are indicated by black arrows, the Aβ42 amino acid sequence is depicted by red letters, light blue letters indicate the unique C-terminal sequence of Aβ175 (which is not present in APP full length protein), light blue underlined letters highlight the amino acids of the unique peptide detected by IP-MS as displayed in D. **C. Western blot analysis of Aβ175 in HEK293 cells.** pCMV-Aβ175-cDNA indicates linear, i.e. canonical mRNA-driven expression of Aβ175 cDNA. Mock transfection with empty vector (pCircRNA-DMo) was included as control. circAβ-a indicates expression after pCircRNA-DMo-Aβ-a transfections in HEK293 cells. The peptides detected are indicated on the right, Aβ175 marks the migration of the circAβ-a-derived protein, β-actin served as the loading control. SeeBlue™ Plus2 pre-stained protein standard was utilized as a protein size marker. * represents possible protein products that are the result of alternative translational initiation at AUG codons further downstream. **D. Mass spectrometry to demonstrate circular circAβ-a translation.** The unique peptide (see B for details) was enriched via immunoprecipitation of Aβ175 with the anti-Aβ antibody in HEK293 cells transfected with the pCircRNA-DMo-Aβ-a expression vector (4G8 and 6E10), the scale of the left ordinate is the relative signal intensity, absolute intensities are displayed by the ordinate to the right, the x-axis denotes m/z values. **E. FLAG-tagged circAβ-a (circAβ-a-FLAG).** The FLAG peptide tag, i.e. DYKDDDDKPP, was fused to the C-terminus of the circAβ-a ORF. **F. Western blot analysis for Aβ175-FLAG expression in N2a cells with the** anti-FLAG antibody (M2). Control represents empty vector (pCircRNA-DMo) mock transfections, circAβ-a-FLAG displays expression of the pCircRNA-DMo-Aβ-a-FLAG vector, Precision plus protein™ dual colour Standards served as size marker and β-actin as the loading control. **G. Mass spectrometry of the unique peptide representing a circular translation of circAβ-a-FLAG.** This peptide was enriched via immunoprecipitation of Aβ175 with anti-Aβ antibody (4G8 and 6E10), the scale of the left ordinate is the relative signal intensity, absolute intensities are displayed by the right ordinate, the x-axis denotes m/z values.

Due to the fact that the 524 nt of circAβ-a are not divisible by three, the translation product is not completely similar to the corresponding APP segment, due to a unique 17 amino acids (aa) extension, contributing to the C-terminus of the Aβ175 protein. These aa residues are a consequence of circular translation into a different reading frame at the junction of exon 17 and 14, contributed by the latter exon out of frame. This aa sequence is absent from full-length APP (Fig. 2A, Supplementary data-4). To provide further experimental evidence for circAβ-a translation, we performed mass spectrometry analysis to verify Aβ175 specific peptides. For enrichment, Aβ175 was immunoprecipitated with anti-Aβ antibodies (6E10, 4G8) from lysates of circAβ-a overexpressing HEK293 cells. Subsequent mass spectrometry revealed a peptide (SKTIOMTSWPT) corresponding to part of the unique C-terminal portion of Aβ175 (Fig. 2C and D). Therefore, this analysis provides strong evidence for circAβ-a translation in the HEK293 transfection assays. Finally, we modified the C-terminal portion of Aβ175 in circAβ-a ORF adding the FLAG-tag sequence (DYKDDDDKPP). The resulting recombinant expression vector was designated as pCircRNA-DMo-Aβ-a-FLAG (Fig. 2E). To validate circAβ-a-FLAG translation in neuronal cell lines, we utilized transient transfection assays in N2a cells. Western blot analysis with an anti-FLAG monoclonal antibody (M2) revealed a specific signal of approximately 25 kDa (Fig. 2F) confirming that circAβ-a-FLAG also was an efficient template for protein biosynthesis in N2a cells. Similar to our HEK 293 cell system (see above) we resorted to mass spectrometry to further validate the Western blot results. The FLAG-tagged protein (Aβ175-FLAG) was enriched via immunoprecipitation with anti-Aβ antibodies (6E10, 4G8) in circAβ-a-FLAG overexpressing N2a cells.

Subsequent mass spectrometry (MS) analysis, identified a peptide, which is a specific product of circAβ-a-FLAG translation; in fact, it represented the C-terminal fusion of Aβ175 and the added FLAG tag (TIOMTSWPTDYKDDDDKPP, Fig. G). This finding further confirmed protein translation of circAβ-a-FLAG in living cells.

### Aβ175 generates Aβ peptides in HEK293 cells

Translation of the Aβ-related protein from circAβ-a RNA templates in our assays suggested putative functions of the circRNA in the biogenesis of Aβ peptides. The amino acid sequence of Aβ175 contains β and γ-secretase cleavage sites. This implied that amyloid beta peptides might be generated from Aβ175 *via* β- and γ-secretase-mediated cleavage (Fig. 2B). For circAβ-a overexpressing HEK293 cells, we utilized immunoprecipitation coupled to Western blotting (IP-WB) with anti-Aβ-antibodies (for IP: 6E10, 4G8, mouse antibody) to test for Aβ peptides in conditioned cell culture medium (CM). We obtained a specific signal corresponding to Aβ by Western blot analysis (i.e. after IP) with the Aβ antibody D54D2 (rabbit antibody). Compared to HEK293 mock transfections with empty vector, which served as negative control, pCircRNA-DMo-Aβ-a enhanced the expression of the Aβ peptide by more than 4.4-fold (Fig. 3A and B, Supplementary data-5). These results confirmed an alternative route to Aβ biosynthesis via circAβ-a and its translational product Aβ175 (Fig. 3A and B).

**Fig. 3.**
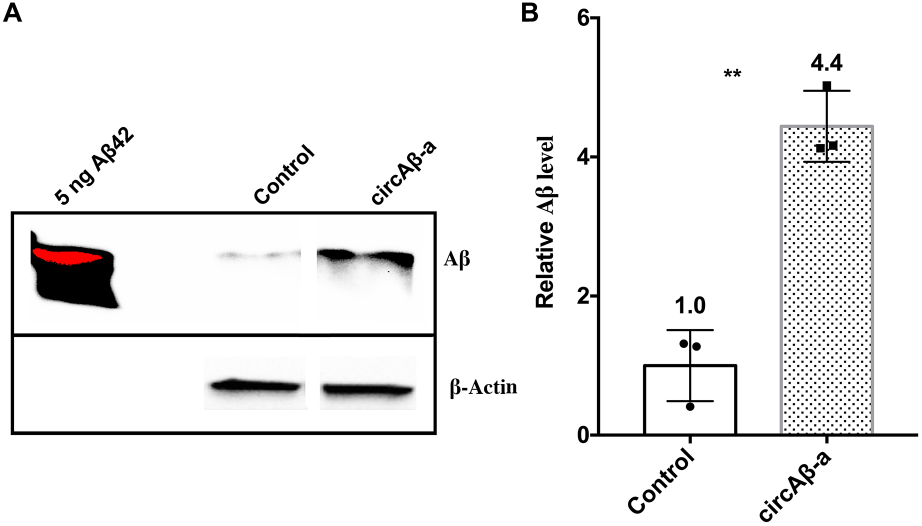
circAβ-a overexpression generates Aβ peptides. **A. IP-WB of Aβ peptides in the conditioned medium of circAβ-a overexpressing cells**, conditioned cell culture medium for HEK293 cells, transfected with the circAβ-a overexpression vector was utilized for immunoprecipitation with antibodies against Aβ (6E10, 4G8; mouse antibodies). Control represents the IP-WB results for mock transfections (pCircRNA-DMo), circAβ-a indicates pCircRNA-DMo-Aβ-a transfections, rabbit D54D2 antibody specific for Aβ was utilized in this Western blot analysis, β-actin served as loading control and 5 ng of *in vitro* synthesised Aβ42 served as Aβ migration maker. The red colour for the Aβ42 signal in the left lane was the result of over-exposure. **B. Quantification of A.** Student’s T-tests were performed against the control sample; **, P ≤ 0.01; n = 3. Details for the three replicates of IP-WB results are provided in Supplementary data-5.

## DISCUSSION

The biogenesis of Aβ in familial AD is well-documented, but the Aβ production in case of sporadic AD remains elusive [1,40–42]. Here, we report the identification of circAβ-a and the analysis of its formation in AD patients, non-dementia controls, and transient transfection assays [63]. With the aid of intron-mediated enhancement, we confirmed that the circRNA harbors a functional ORF and serves as template for the biosynthesis of an Aβ-related protein (Aβ175) [55,63]. In addition, Aβ175 is being processed into Aβ peptides, representing a novel route of Aβ generation that might give rise to new perspectives on the molecular mechanisms leading to the manifestation of Alzheimer’s disease (Fig. 4) [63]. The portrayed mechanism is substantially different from the ‘conventional’ Aβ accumulation caused by the pathophysiological dysregulation of full-length APP proteolytic processing (Fig. 4) [14]. In particular, unlike specific mutations in the APP gene, which are known to be commonly responsible for familial forms of AD[14], no such specific mutations are required for the biogenesis of circAβ-a. circRNA biogenesis is largely directed by the complementary elements in surrounding intronic regions [64]. This implies that maybe even the entire human population expresses circAβ-a RNA, indicating that it might play key roles in the pathogenesis of sporadic AD. Our data describe potential functions for circAβ-a and reveal a novel mechanism for Aβ-related protein biosynthesis. Already, it had been demonstrated that some circRNA translation could be activated under certain stress conditions [56,65,66]. Further analysis is required for the investigation whether circAβ-a translation is elevated in the ageing brain, which occurs under various cellular stresses, such as oxidative damage[67–69]. Ultimately, it is essential to investigate the ratio of Aβ peptides as a result of circAβ-a translation compared to APP full-length protein. Furthermore, compared to full-length APP protein, Aβ175 lacks an N-terminal domain, has a shorter and modified C-terminus and presumably a different tertiary structure. Consequently, it will be important to investigate whether a different structure exposes the corresponding amino acid residues to increase processing of Aβ175 by secretases, thus generating more Aβ peptides, especially in the context of the ageing human brain and during neurodegenerative diseases.

**Fig. 4.**
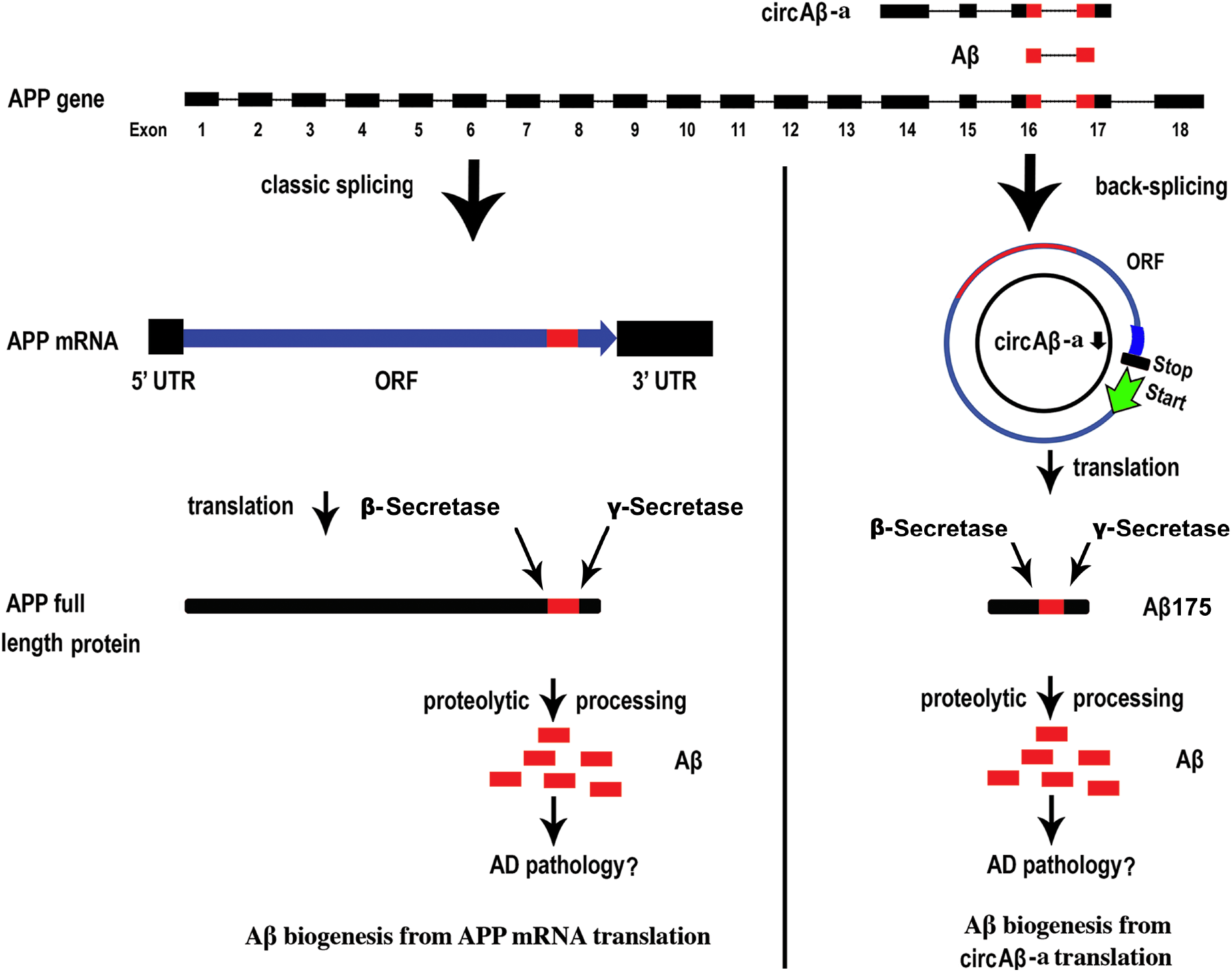
The alternative pathway of Aβ biogenesis in Alzheimer’s disease. At the top, exon sequences containing circAβ-a and Aβ peptides (in red) are aligned with a scheme of the full-length APP gene (not drawn to scale). On the left, linear APP mRNA transcribed from the APP gene locus undergoes the canonical splicing pathway before being translated into full-length APP protein. Proteolytic processing of APP protein generates Aβ peptides (e.g., Aβ40, Aβ42, in red colour), which may play causative roles in the AD pathology. On the right, circAβ-a is synthesised by back-splicing of the APP gene. The open reading frame (ORF) is in blue, with the Aβ sequence in red. The presumed translational start codon is depicted by a green arrow and the stop codon by a black bar. Translation of circAβ-a produces Aβ-related peptide (Aβ175), which is further processed to form Aβ.

In summary, the investigation of biological roles of circAβ-a will lead to new perspectives in the search of a mechanism underlying Alzheimer’s disease and provides routes to novel strategies for the design of disease-modifying drugs[54]. For example, an antisense oligonucleotide annealing to the junction region of circAβ-a could specifically reduce the circRNA expression without affecting the cognate APP mRNA (Dingding Mo, unpublished data), representing a promise of next generation of Alzheimer’s disease therapy [70].

## METHODS

### circAβ-a identification via RT-PCR and sequencing

The APP (amyloid beta precursor protein) gene-derived circRNAs were amplified *via* RT-PCR with specific ‘divergent’, i.e., head-to-head oriented primers targeting protein coding exon 17 of the APP gene. One μg of total RNA samples of the human frontal cortex of Alzheimer’s disease individuals and health control individuals (from the London Neurodegenerative Diseases Brain Bank, London, UK; the human brain samples study was approved by the ethical committees of Leuven University and UZ Leuven.) were used as template. cDNA synthesis was performed with SuperScript™ II Reverse Transcriptase (18064022, Invitrogen) with random hexamers primer, according to the manufacturer’s recommendations. PCR was performed with Phusion^®^ High-Fidelity DNA Polymerase (M0530L, NEB) for 40 cycles with the protocol provided by the manufacturer.

For RNase R treatment, about 15 μg of total RNA of human frontal cortex were treated with 10 units of RNase R (RNR07250, Epicentre) for 1 hour at 37 °C and purified by phenolchloroform extraction. 250 ng of the resulting RNA samples were utilized for subsequent cDNA synthesis and PCR amplifications. PCR products were purified by PCR purification kit (28104, QIAGEN) and subsequently verified by Sanger sequencing.

### Plasmid construction

Human hsa_circ_0007556 (circBase[60]), is referred to as circAβ-a in this study. The exon DNA representing circAβ-a (exon 14, 15, 16, 17 of APP gene, GRCh37/hg19, chr21:27264033-27284274) was inserted into pCircRNA-DMo vectors to generate pCircRNA-DMo-Aβ-a as described previously[55]. As positive control for Aβ175 protein expression, the cDNA containing its ORF was inserted into the pCMV-MIR vector (OriGene) to result in pCMV-Aβ175-cDNA. A FLAG tag sequence (DYKDDDDKPP) was added to pCircRNA-DMo-Aβ-a to generate pCircRNA-DMo-Aβ-a-FLAG. Recombinant plasmids were purified with EndoFree Plasmid Maxi Kit (12362, QIAGEN). Oligonucleotide sequences and further details are provided in Supplementary Table 1. All plasmids were verified by restriction endonuclease digestions and Sanger sequencing.

### Cell culture and plasmid DNA transfection

Cell culture and plasmid DNA transfection in HEK293 or N2a cells were performed as previously described[55]. In addition, 50 μM α-secretase ADAM10 inhibitor GI254023X (SML0789 Sigma), 10 μM β-Secretase Inhibitor IV-CAS 797035-11-1-Calbiochem (565788-1MG, Millipore) and 50 μM γ-Secretase Inhibitor (Begacestat, PZ0187, Sigma) (SCP0004, Sigma) were added to the cell culture for 24 hours.

### Total RNA isolation and qRT-qPCR

Total RNA from HEK293 cells and human brain prefrontal cortex were isolated with the TRIzol reagent (Ambion) according to the manufacturer’s recommendations. cDNA synthesis and qRT-PCR were performed as previously described[55]. Details of qRT-PCR oligonucleotides are provided in Supplementary Table 1.

### Northern blot

Northern blot hybridizations were conducted with the NorthernMax™ Kit (AM1940, Ambion) as previously described[55]. In brief, 15 μg total RNA from HEK293 cells were separated on 5% Criterion™ TBE polyacrylamide gels (#3450048, Bio-Rad) and transferred to positively charged nylon membrane. Hybridization was performed with a 5’ P^32^-labeled DNA oligonucleotide for overnight at 42 °C (NB-R1: CCCACCATGAGTCCAATGATTGCACCTTTGTTTGAACCCACAT CTTCTGCAAAGAACACC). Membranes were washed at 42 °C with the kit provided buffer according the manufacturer’s recommendations. For RNase R treatment, 15 μg total RNA were digested with 10 units of RNase R (RNR07250, Epicentre) for 1 hour at 37 °C; RNAs were separated by gel electrophoresis and analysed by Northern blot as above described.

### Western blot analysis

Protein lysates were prepared with RIPA Buffer (50 mM Tris-HCl pH 8.0, 150 mM NaCl, 1% (v/v) NP40, 0.1% (w/v) SDS, 0.5% (w/v) Na-deoxycholate, Roche complete Protease Inhibitor and PhosSTOP Phosphatase Inhibitor). Aliquots representing 40 μg total proteins were separated on Novex™ 4-20% Tris-Glycine Mini Gels (Invitrogen) or 4-20% Criterion™ TGX Stain-Free™ Protein gels (Bio-Rad) and transferred to 0.2 μm nitrocellulose membranes (10600002, Amersham). Immunoblotting was performed with anti-β-amyloid antibody (clone WO2, MABN10, Sigma-Aldrich) (β-Amyloid [D54D2] XP^®^ Rabbit mAb #8243, Cell signalling Technology), monoclonal ANTI-FLAG M2 antibody (F3165, Sigma-Aldrich) and anti-β-actin (#A5441, Sigma) antibody as previously described[55]. Aβ42 peptide (A9810, Sigma) was prepared in DMSO. Quantitative analyses were performed with ImageJ (NIH).

### Immunoprecipitation–Mass Spectrometry (IP-MS) of circAβ-a derived protein

pCircRNA-DMo-Aβ-a or pCircRNA-DMo-Aβ-a-FLAG was transfected in HEK293 or N2a cells with secretase inhibitors for 24 hours as previously described. Cells were collected and lysed in RIPA buffer[55]. Immunoprecipitations were performed with anti-β-Amyloid antibodies 6E10 and 4G8 (803001, 800701, BioLegend Inc.) bound to Dynabeads™ Protein A, G (10002D, 10004D, Invitrogen). Immunoprecipitated proteins were digested on beads by trypsin (V5280, Omega). Mass spectrometry was performed as previously described[55].

### Immunoprecipitation/Western blotting (IP-WB) of Aβ peptides

Aβ peptide detection was performed through immunoprecipitation of conditioned medium (CM), followed by Western blot analysis as previously described[71]. In brief, HEK293 cells transfected with pCircRNA-DMo-Aβ-a or empty vector (pCircRNA-DMo) were cultured in serum free medium overnight. Then CM was prepared with protease and phosphatase inhibitors (Roche) and pre-cleaned with protein A/G beads (Dynabeads™ Protein A, 10002D, Dynabeads™ Protein G, 10004D, Invitrogen). Immunoprecipitation was conducted with a mixture of Aβ antibodies (6E10, 4G8, BioLegend Inc.) and protein A/G beads. Precipitated peptides were subsequently dissolved in SDS loading buffer and analysed by Western blot with an antibody derived against Aβ (D54D2, Cell signalling Technology).

## Supporting information

Suppelmentary data-1

Suppelmentary data-2

Suppelmentary data-3

Suppelmentary data-4

Suppelmentary data-5

Suppelmentary table-1

## Acknowledgements

The authors acknowledge the lab led by Prof. Linda Partridge at the Max Planck Institute for Biology of Ageing (MPI-AGE) for sharing chemicals and instruments. The author appreciates Dr. Gabriella B Lundkvist for support during the study. The authors acknowledge the support from the lab of Prof. Bart De Strooper at the VIB-KU Leuven Center for Brain & Disease Research. The authors also thank Stephanie Klco-Brosius for editorial advice.

## Author Contributions

D.M. designed and conceived the study. D.M. performed the experiments, analysed the results, prepared the figures and wrote the manuscript. X.L. performed the mass spectrometry analysis. X.L., C.R., T.R., B.V.S. and J.B. participated in project discussions, data analysis, results integration, and manuscript revising. J.B. prepared the supplementary data-4A.

## Supplementary Materials

Supplementary data1, 2, 3, 4

Supplementary table-1

## Data and materials availability

For correspondence, please contact Dr. Dingding Mo.

