## Supplementary material for "The role of Aβ circRNA in Alzheimer’s disease: alternative mechanism of Aβ biogenesis from Aβ circRNA translation": Suppelmentary data-1

```
                                            10        20        30        40        50
                                     CGTCTTGG-CCAACATGATTAGTGAACCAAGGATCAGTTACGGAAACGAT
circAB-a-JR1-amplicon.seq(1>499)  →  CGTCTTGG-CCAACATGATTAGTGAACCAAGGATCAGTTACGGAAACGAT
R-AD3-R-58_1.seq(8>437)           ←               CATGATTAGTGAACCAAGGATCAGTTACGGAAACGAT
R-ND3-R-59_1.seq(17>460)          ←              ACATGATTAGTGAACCAAGGATCAGTTACGGAAACGAT
AD1-R-48_1.seq(31>437)            ←              ACATGATTAGTGAACCAAGGATCAGTTACGGAAACGAT
R-ND3-F-50_1.seq(9>455)           →                                              AACGAT
```

```
                                            60        70        80        90       100
                                     GCTCTCATGC-CAT-CTTTGACCGAAACGAAAAC-CACCGTGGAGCTCCT
circAB-a-JR1-amplicon.seq(1>499)  →  GCTCTCATGC-CAT-CTTTGACCGAAACGAAAAC-CACCGTGGAGCTCCT
R-AD3-R-58_1.seq(8>437)           ←  GCTCTCATGC-CAT-CTTTGACCGAAACGAAAAC-CACCGTGGAGCTCCT
R-ND3-R-59_1.seq(17>460)          ←  GCTCTCATGC-CAT-CTTTGACCGAAACGAAAAC-CACCGTGGAGCTCCT
AD1-R-48_1.seq(31>437)            ←  GCTCTCATGC-CAT-CTTTGACCGAAACGAAAAC-CACCGTGGAGCTCCT
R-ND3-F-50_1.seq(9>455)           →  GCTCTCATGC-CAT-CTTTGACCGAAACGAAAAC-CACCGTGGAGCTCCT
R-AD3-F-49_1.seq(9>440)           →             CAT-CTTTGACCGAAACGAAAAC-CACCGTGGAGCTCCT
AD1-F-45_1.seq(29>436)            →                                    CACCGTGGAGCTCCT
```

```
                                           110       120       130       140       150
                                     TCCCGTGAATGGAGAGTTCAGCCTGGACGATCTCCAGCCGTGGCATTCTT
circAB-a-JR1-amplicon.seq(1>499)  →  TCCCGTGAATGGAGAGTTCAGCCTGGACGATCTCCAGCCGTGGCATTCTT
R-AD3-R-58_1.seq(8>437)           ←  TCCCGTGAATGGAGAGTTCAGCCTGGACGATCTCCAGCCGTGGCATTCTT
R-ND3-R-59_1.seq(17>460)          ←  TCCCGTGAATGGAGAGTTCAGCCTGGACGATCTCCAGCCGTGGCATTCTT
AD1-R-48_1.seq(31>437)            ←  TCCCGTGAATGGAGAGTTCAGCCTGGACGATCTCCAGCCGTGGCATTCTT
R-ND3-F-50_1.seq(9>455)           →  TCCCGTGAATGGAGAGTTCAGCCTGGACGATCTCCAGCCGTGGCATTCTT
R-AD3-F-49_1.seq(9>440)           →  TCCCGTGAATGGAGAGTTCAGCCTGGACGATCTCCAGCCGTGGCATTCTT
AD1-F-45_1.seq(29>436)            →  TCCCGTGAATGGAGAGTTCAGCCTGGACGATCTCCAGCCGTGGCATTCTT
```

```
                                           160       170       180       190       200
                                     TTGGGGCTGACTCTGTGCCAGCCAACACAGAAAACGAAGTTGAGCCTGTT
circAB-a-JR1-amplicon.seq(1>499)  →  TTGGGGCTGACTCTGTGCCAGCCAACACAGAAAACGAAGTTGAGCCTGTT
R-AD3-R-58_1.seq(8>437)           ←  TTGGGGCTGACTCTGTGCCAGCCAACACAGAAAACGAAGTTGAGCCTGTT
R-ND3-R-59_1.seq(17>460)          ←  TTGGGGCTGACTCTGTGCCAGCCAACACAGAAAACGAAGTTGAGCCTGTT
AD1-R-48_1.seq(31>437)            ←  TTGGGGCTGACTCTGTGCCAGCCAACACAGAAAACGAAGTTGAGCCTGTT
R-ND3-F-50_1.seq(9>455)           →  TTGGGGCTGACTCTGTGCCAGCCAACACAGAAAACGAAGTTGAGCCTGTT
R-AD3-F-49_1.seq(9>440)           →  TTGGGGCTGACTCTGTGCCAGCCAACACAGAAAACGAAGTTGAGCCTGTT
AD1-F-45_1.seq(29>436)            →  TTGGGGCTGACTCTGTGCCAGCCAACACAGAAAACGAAGTTGAGCCTGTT
```

```
                                           210       220       230       240       250
                                     GATGCCCGCCCTGCTGCCGACCGAGGACTGACCACTCGACCAGGTTCTGG
circAB-a-JR1-amplicon.seq(1>499)  →  GATGCCCGCCCTGCTGCCGACCGAGGACTGACCACTCGACCAGGTTCTGG
R-AD3-R-58_1.seq(8>437)           ←  GATGCCCGCCCTGCTGCCGACCGAGGACTGACCACTCGACCAGGTTCTGG
R-ND3-R-59_1.seq(17>460)          ←  GATGCCCGCCCTGCTGCCGACCGAGGACTGACCACTCGACCAGGTTCTGG
AD1-R-48_1.seq(31>437)            ←  GATGCCCGCCCTGCTGCCGACCGAGGACTGACCACTCGACCAGGTTCTGG
R-ND3-F-50_1.seq(9>455)           →  GATGCCCGCCCTGCTGCCGACCGAGGACTGACCACTCGACCAGGTTCTGG
R-AD3-F-49_1.seq(9>440)           →  GATGCCCGCCCTGCTGCCGACCGAGGACTGACCACTCGACCAGGTTCTGG
AD1-F-45_1.seq(29>436)            →  GATGCCCGCCCTGCTGCCGACCGAGGACTGACCACTCGACCAGGTTCTGG
```

```
                                           260       270       280       290       300
                                     GTTGACAAATATCAAGACGGAGGAGATCTCTGAAGTGAAGATGGATGCAG
circAB-a-JR1-amplicon.seq(1>499)  →  GTTGACAAATATCAAGACGGAGGAGATCTCTGAAGTGAAGATGGATGCAG
R-AD3-R-58_1.seq(8>437)           ←  GTTGACAAATATCAAGACGGAGGAGATCTCTGAAGTGAAGATGGATGCAG
R-ND3-R-59_1.seq(17>460)          ←  GTTGACAAATATCAAGACGGAGGAGATCTCTGAAGTGAAGATGGATGCAG
AD1-R-48_1.seq(31>437)            ←  GTTGACAAATATCAAGACGGAGGAGATCTCTGAAGTGAAGATGGATGCAG
R-ND3-F-50_1.seq(9>455)           →  GTTGACAAATATCAAGACGGAGGAGATCTCTGAAGTGAAGATGGATGCAG
R-AD3-F-49_1.seq(9>440)           →  GTTGACAAATATCAAGACGGAGGAGATCTCTGAAGTGAAGATGGATGCAG
AD1-F-45_1.seq(29>436)            →  GTTGACAAATATCAAGACGGAGGAGATCTCTGAAGTGAAGATGGATGCAG
```

```
                                         310       320       330       340       350
                                AATTCCGACATGACTCAGGATATGAAGTTCATCATCAAAAATTGGTGTTC
circAB-a-JR1-amplicon.seq(1>499)  →  AATTCCGACATGACTCAGGATATGAAGTTCATCATCAAAAATTGGTGTTC
R-AD3-R-58_1.seq(8>437)           ←  AATTCCGACATGACTCAGGATATGAAGTTCATCATCAAAAATTGGTGTTC
R-ND3-R-59_1.seq(17>460)          ←  AATTCCGACATGACTCAGGATATGAAGTTCATCATCAAAAATTGGTGTTC
AD1-R-48_1.seq(31>437)            ←  AATTCCGACATGACTCAGGATATGAAGTTCATCATCAAAAATTGGTGTTC
R-ND3-F-50_1.seq(9>455)           →  AATTCCGACATGACTCAGGATATGAAGTTCATCATCAAAAATTGGTGTTC
R-AD3-F-49_1.seq(9>440)           →  AATTCCGACATGACTCAGGATATGAAGTTCATCATCAAAAATTGGTGTTC
AD1-F-45_1.seq(29>436)            →  AATTCCGACATGACTCAGGATATGAAGTTCATCATCAAAAATTGGTGTTC
```

```
                                         360       370       380       390       400
                                TTTGCAGAAGATGTGGGTTCAAACAAAGGTGCAATCATTGGACTCATGGT
circAB-a-JR1-amplicon.seq(1>499)  →  TTTGCAGAAGATGTGGGTTCAAACAAAGGTGCAATCATTGGACTCATGGT
R-AD3-R-58_1.seq(8>437)           ←  TTTGCAGAAGATGTGGGTTCAAACAAAGGTGCAATCATTGGACTCATGGT
R-ND3-R-59_1.seq(17>460)          ←  TTTGCAGAAGATGTGGGTTCAAACAAAGGTGCAATCATTGGACTCATGGT
AD1-R-48_1.seq(31>437)            ←  TTTGCAGAAGATGTGGGTTCAAACAAAGGTGCAATCATTGGACTCATGGT
R-ND3-F-50_1.seq(9>455)           →  TTTGCAGAAGATGTGGGTTCAAACAAAGGTGCAATCATTGGACTCATGGT
R-AD3-F-49_1.seq(9>440)           →  TTTGCAGAAGATGTGGGTTCAAACAAAGGTGCAATCATTGGACTCATGGT
AD1-F-45_1.seq(29>436)            →  TTTGCAGAAGATGTGGGTTCAAACAAAGGTGCAATCATTGGACTCATGGT
```

```
                                         410       420       430       440       450
                                GGGCGGTGTTGTCATAGCGACA-GTGATCGTCATCACCTTGGTGATG-CT
circAB-a-JR1-amplicon.seq(1>499)  →  GGGCGGTGTTGTCATAGCGACA-GTGATCGTCATCACCTTGGTGATG-CT
R-AD3-R-58_1.seq(8>437)           ←  GGGCGGTGTTGTCATAGCGACA-GTGATCGTCATCACCTTGGTGATG
R-ND3-R-59_1.seq(17>460)          ←  GGGCGGTGTTGTCATAGCGACA-GTGATCGTCATCACCTTGGTGATG-CT
AD1-R-48_1.seq(31>437)            ←  GGGCGGTGTTGTCATAGCGACA
R-ND3-F-50_1.seq(9>455)           →  GGGCGGTGTTGTCATAGCGACA-GTGATCGTCATCACCTTGGTGATG-CT
R-AD3-F-49_1.seq(9>440)           →  GGGCGGTGTTGTCATAGCGACA-GTGATCGTCATCACCTTGGTGATG-CT
AD1-F-45_1.seq(29>436)            →  GGGCGGTGTTGTCATAGCGACA-GTGATCGTCATCACCTTGGTGATG-CT
```

```
                                         460       470       480       490       500
                                GAAGAAGAAAC-AGTACACATCCATTCATCATGGTGTGGTGGAGATGAGC
circAB-a-JR1-amplicon.seq(1>499)  →  GAAGAAGAAAC-AGTACACATCCATTCATCATGGTGTGGTGGAGATGAGC
R-ND3-R-59_1.seq(17>460)          ←  GAAGAAGAAAC
R-ND3-F-50_1.seq(9>455)           →  GAAGAAGAAAC-AGTACACATCCATTCATCATGGTGTGGTGGAGATG
R-AD3-F-49_1.seq(9>440)           →  GAAGAAGAAAC-AGTACACATCCATTCATCATGGTGTGGTGGAGATGA
AD1-F-45_1.seq(29>436)            →  GAAGAAGAAAC-AGTACACATCCATTCATCATGGTGTGGTGGAGAT
```

```
                                         510
                                TGCTTC
circAB-a-JR1-amplicon.seq(1>499)  →  TGCTTC
```
