## Supplementary material for "The role of Aβ circRNA in Alzheimer’s disease: alternative mechanism of Aβ biogenesis from Aβ circRNA translation": Suppelmentary data-2

```
                                                                10        20        30        40
                                                   TTGTCATAGCGACAGTGATCGTCATCACCTTGGTGATGCTGAAG

AD3-R-60_1.seq(7>104)                     ←        TTGTCATAGCGACAGTGATCGTCATCACCTTGGTGATGCTGAAG
ND3-R-61_1.seq(4>98)                      ←           TCATAGCGACAGTGATCGTCATCACCTTGGTGATGCTGAAG
circAB-a-junction  region-WT.seq(1>148)   →          GTCATAGCGACAGTGATCGTCATCACCTTGGTGATGCTGAAG

                                                        50        60        70        80
                                                   AAGAAACAGTACACATCCATTCATCATGGTGTGGTGGAGATGAG

AD3-R-60_1.seq(7>104)                     ←        AAGAAACAGTACACATCCATTCATCATGGTGTGGTGGAGATGAG
ND3-R-61_1.seq(4>98)                      ←        AAGAAACAGTACACATCCATTCATCATGGTGTGGTGGAGATGAG
circAB-a-junction  region-WT.seq(1>148)   →        AAGAAACAGTACACATCCATTCATCATGGTGTGGTGGAGATGAG
ND3-F-52_1.seq(14>113)                    →                GTACACATCCATTCATCATGGTGTGGTGGAGATGAG
AD3-F-51_1.seq(3>107)                     →                GTACACATCCATTCATCATGGTGTGGTGGAGATGAG

                                                    90       100       110       120       130
                                                   CTGCTTCAGAAAGAGCAAAACTATTCAGATGACGTCTTGGCCAA

AD3-R-60_1.seq(7>104)                     ←        CTGCTTCAGA
ND3-R-61_1.seq(4>98)                      ←        CTGCTTCAGA
circAB-a-junction  region-WT.seq(1>148)   →        CTGCTTCAGAAAGAGCAAAACTATTCAGATGACGTCTTGGCCAA
ND3-F-52_1.seq(14>113)                    →        CTGCTTCAGAAAGAGCAAAACTATTCAGATGACGTCTTGGCCAA
AD3-F-51_1.seq(3>107)                     →        CTGCTTCAGAAAGAGCAAAACTATTCAGATGACGTCTTGGCCAA

                                                         140       150
                                                   CATGATTAGTGAACCAAGAAGCTTG

circAB-a-junction  region-WT.seq(1>148)   →        CATGATTAGTGAACCAAG
ND3-F-52_1.seq(14>113)                    →        CATGATTAGTGAACCAAGAA
AD3-F-51_1.seq(3>107)                     →        CATGATTAGTGAACCAAGAAGCTTG
```
