## Supplementary material for "The role of Aβ circRNA in Alzheimer’s disease: alternative mechanism of Aβ biogenesis from Aβ circRNA translation": Suppelmentary data-3

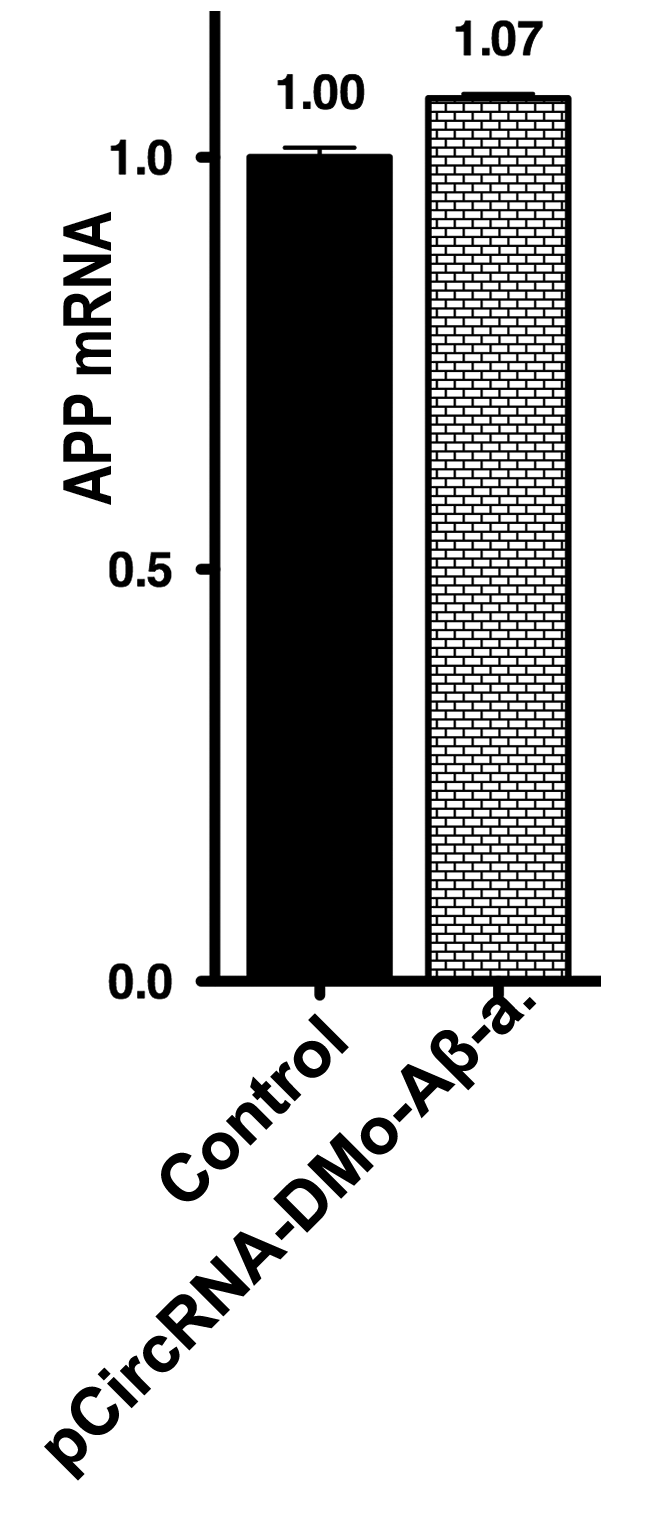

**Supplementary data-3. Endogenous APP mRNA expression in HEK293 cells overexpressing circAβ-a.**

qRT-PCR analysis of endogenous APP mRNA expression in circAβ-a overexpressing HEK293 cells. Primers targeted exons 3 and 4, which are not contained in circAβ-a: Control, pCircRNA-DMo empty vector, pCircRNA-DMo-Aβ-a. No significant differences between different groups were observed. n = 4.
