## Supplementary material for "The role of Aβ circRNA in Alzheimer’s disease: alternative mechanism of Aβ biogenesis from Aβ circRNA translation": Suppelmentary data-4

**Supplementary data-4**

**A. Open reading frame of circAβ-a**

```
5' end of exon 14
AUGAGCUGCUUCAGAAAGAGCAAAACUAUUCAGAUGACGUCUUGGCCAAC


AUGAUUAGUGAACCAAGGAUCAGUUACGGAAACGAUGCUCUCAUGCCAUC
-M--I--S--E--P--R--I--S--Y--G--N--D--A--L--M--P--S
 1                                                         17

UUUGACCGAAACGAAAACCACCGUGGAGCUCCUUCCCGUGAAUGGAGAGU
--L--T--E--T--K--T--T--V--E--L--L--P--V--N--G--E
  18                                                    33

UCAGCCUGGACGAUCUCCAGCCGUGGCAUUCUUUUGGGGCUGACUCUGUG
F--S--L--D--D--L--Q--P--W--H--S--F--G--A--D--S--V
34                                                       50

                          5' end of exon 15
CCAGCCAACACAGAAAACGAAGUUGAGCCUGUUGAUGCCCGCCCUGCUGC
-P--A--N--T--E--N--E--V--E--P--V--D--A--R--P--A--A
 51                                                       67

                               5' end of exon 16
CGACCGAGGACUGACCACUCGACCAGGUUCUGGGUUGACAAAUAUCAAGA
--D--R--G--L--T--T--R--P--G--S--G--L--T--N--I--K--
  68                                                    83

CGGAGGAGAUCUCUGAAGUGAAGAUGGAUGCAGAAUUCCGACAUGACUCA
T--E--E--I--S--E--V--K--M--D--A--E--F--R--H--D--S
84                                                       100

                                5' end of exon 17
GGAUAUGAAGUUCAUCAUCAAAAAUUGGUGUUCUUUGCAGAAGAUGUGGG
-G--Y--E--V--H--H--Q--K--L--V--F--F--A--E--D--V--G
 101                                                     117

UUCAAACAAAGGUGCAAUCAUUGGACUCAUGGUGGGCGGUGUUGUCAUAG
 -S--N--K--G--A--I--I--G--L--M--V--G--G--V--V--I--
  118                                                   133

CGACAGUGAUCGUCAUCACCUUGGUGAUGCUGAAGAAGAAACAGUACACA
A--T--V--I--V--I--T--L--V--M--L--K--K--K--Q--Y--T
134                                                      150

									 5' end of exon 14
UCCAUUCAUCAUGGUGUGGUGGAGAUGAGCUGCUUCAGAAAGAGCAAAAC
-S--I--H--H--G--V--V--E--M--S--C--F--R--K--S--K--T
 151                                                     167

UAUUCAGAUGACGUCUUGGCCAACAUGAUUAGUGAA...
 -I--Q--M--T--S--W--P--T
  168                      175
```

**Translated sequence of circAβ-a.** Shown is slightly more than a full circle of circAβ-a (extended by the repeated relevant 5' portion of exon 14). For better identification of the regions corresponding to the exons, they are highlighted in an alternating fashion in grey (exons 15 and 17) or are left unmodified (exons 14 and 16). The putative AUG start codon and the corresponding methionine residue (M) is shown in bold green lettering and the stop codon in bold dark red. The Aβ42 amino acid sequence is shown in red lettering and the additional C-terminal amino acids of Aβ175 encoded by exon 14 out of frame, due to the circular nature of the RNA template, are shown in blue. These 17 specific amino acids cannot be generated by the canonical APP mRNA. Peptides confirmed by mass spectrometry are underlined here.

>Aβ175, putative sequence

MISEPRISYGNDALMPSLTETKTTVELLPVNGEFSLDDLQPWHSFGADSVPANTENEVEPVDARPAADRGLTTRPGSGLTNIKTEEISEVKM**DAEFRHDSGYEVHHQKLVFFAEDVGSNKGAIIGLMVGGVVIA**TVIVITLVMLKKKQYTSIHHGVVE**MSCFRKSKTIQMTSWPT**

The Aβ42 sequence is shown in red. The 17 additional C-terminal amino acids encoded by the circular RNA are shown in blue.

**B. Alignment of Aβ175 with APP695**

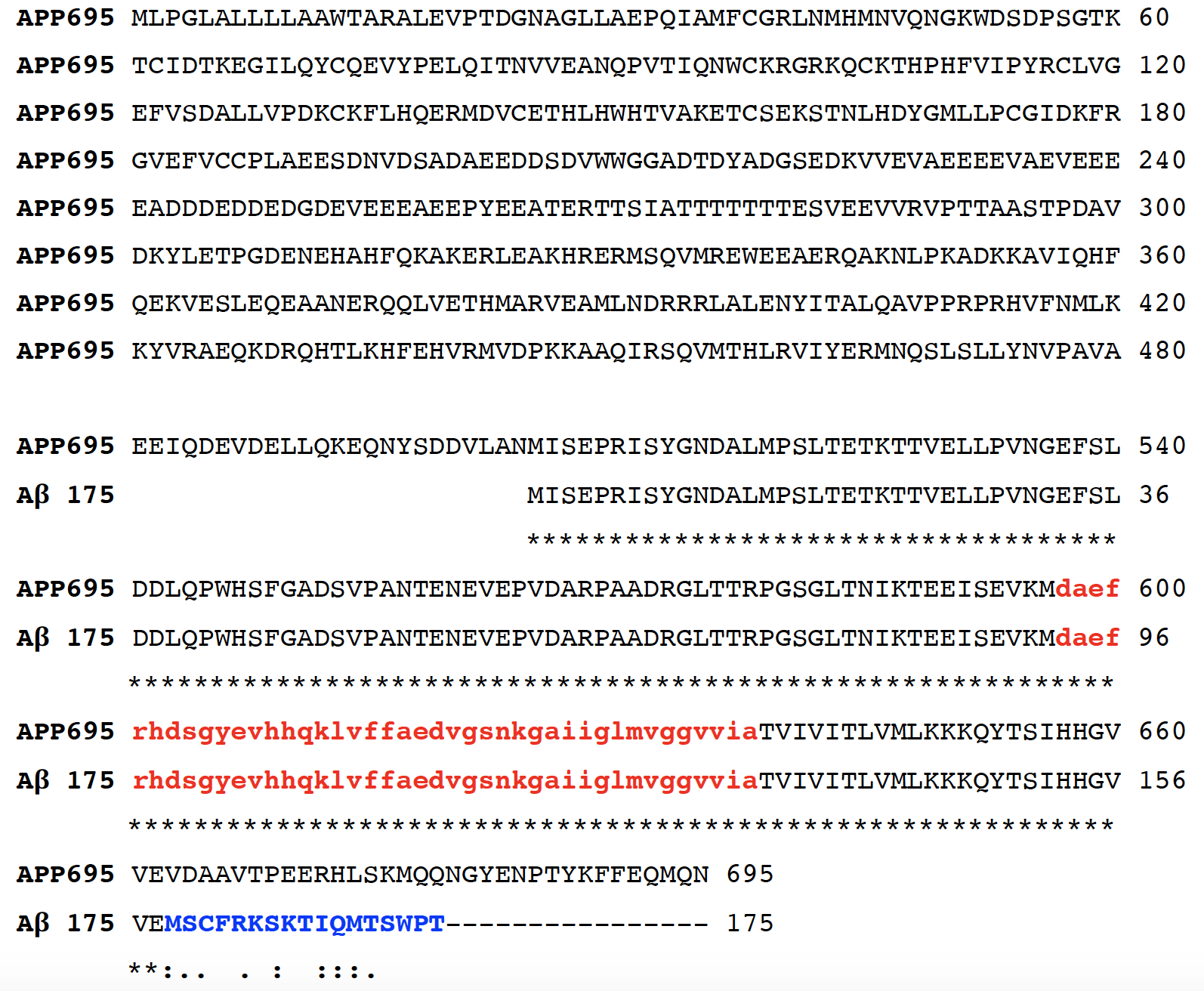

Protein sequence alignment between APP695 and Aβ175 was done through Clustal Omega (<https://www.ebi.ac.uk/Tools/msa/clustalo/>). Aβ42 sequence is shown in red color with lowercase.
