## Supplementary material for "The role of Aβ circRNA in Alzheimer’s disease: alternative mechanism of Aβ biogenesis from Aβ circRNA translation": Suppelmentary data-5

**Supplementary data-5**

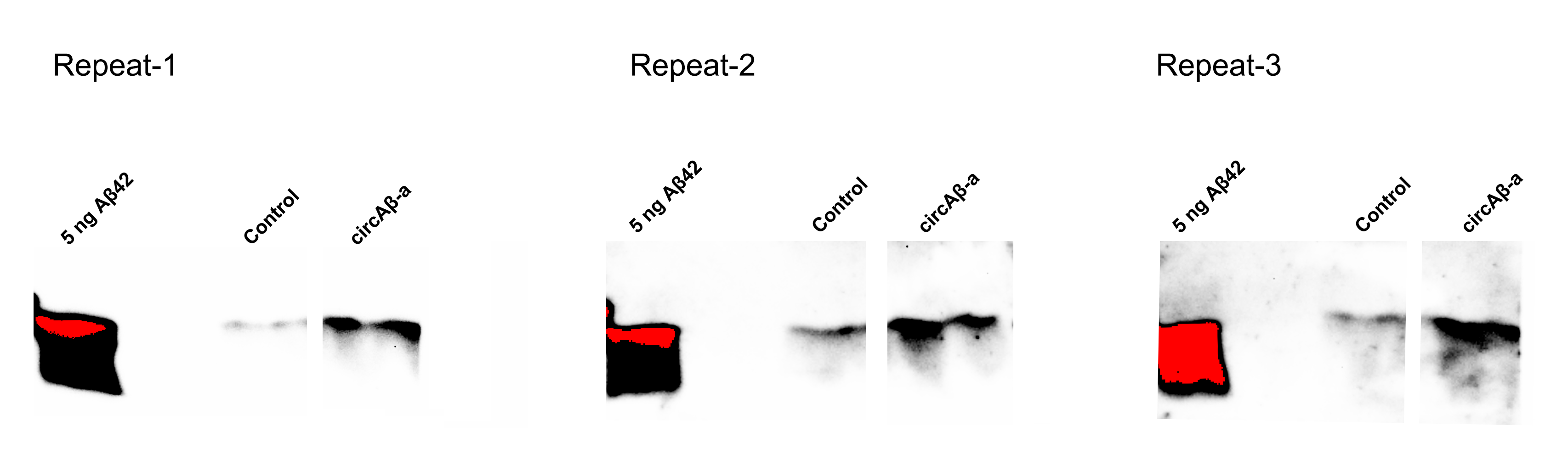

**circAβ-a overexpression generates Aβ peptides.**

**IP-WB of Aβ peptides in the conditioned medium of circAβ-a overexpressing cells**. Conditioned cell culture medium for HEK293 cells, transfected with the circAβ-a overexpression vector was utilized for immunoprecipitation with antibodies against Aβ (6E10, 4G8; mouse antibodies). Control represents the IP-WB results for mock transfections (pCircRNA-DMo), circAβ-a indicates pCircRNA-DMo-Aβ-a transfections, rabbit D54D2 antibody specific for Aβ was utilized in this Western blot analysis, β-Actin served as loading control and 5 ng of *in vitro* synthesised Aβ42 were added as Aβ migration maker. The red colour for the Aβ42 signal in the left most lane was the result of over-exposure. Three repeats are presented here.
