## Supplementary material for "The role of Aβ circRNA in Alzheimer’s disease: alternative mechanism of Aβ biogenesis from Aβ circRNA translation": Suppelmentary table-1

**Supplementary Table 1**

DNA oligonucleotides used in this study

| human circAβ-a expression plasmid construction oligos | Oligonucleotide sequences |
| --- | --- |
| Abeta-circF | gtttgtttttcagATGAGCTGCTTCAGAAAGAGCAAAACT |
| ABeta-circR | gcatggattattacCTCCACCACACCATGATGAATGG |
| circDMO-LF | GTCGACTGGATCcaacgttaaccc |
| DMo-Ab-LR | CTGAAGCAGCTCATctgaaaaacaaacagaatacaacctcagc |
| DMo-Ab-RF | GGTGTGGTGGAGgtaataatccatgcaccgtctcacc |
| circDMO-RR | cactttgCTCGAGctcatcaacatg |
| oligonucleotides for human circAβ-a identification |  |
| circAβ-a-R1 | GAAGCAGCTCATCTCCACCACACC |
| circAβ-a-F1 | CGTCTTGGCCAACATGATTAGTGAACC |
| circAβ-a-F2 | GTCATAGCGACAGTGATCGTC |
| circAβ-a-R2 | CTTGGTTCACTAATCATGTTGGC |
| circAβ-a-F3 | GTGATCGTCATCACCTTGGTGATGC |
| circAβ-a-R3 | CACCATGAGTCCAATGATTGCACC |
| oligonucleotides for human APP mRNA qRT-PCR |  |
| hAPP-mF | TTTGTGATTCCCTACCGCTG |
| hAPP-mR | TGCCAGTGAAGATGAGTTTCG |
| Human ACTB mRNA: |  |
| hACTB-F | ACCTTCTACAATGAGCTGCG |
| hACTB-R | CCTGGATAGCAACGTACATGG |
